# Rescuing Biologically Relevant Consensus Regions Across Replicated Samples

**DOI:** 10.1101/2021.09.23.461528

**Authors:** Vahid Jalili, Marzia Angela Cremona, Fernando Palluzzi

## Abstract

**Motivation:** Protein-DNA binding sites of ChIP-seq experiments are identified where the binding affinity is significant based on a given threshold. The choice of the threshold is a trade-off between conservative region identification and discarding weak, but true binding sites.

**Results:** We argue the biological relevance of weak binding sites and the information they add when *rescued*. The sites are rescued using MSPC, which exploits replicates to lower the threshold required to identify a binding site while keeping a low false-positive rate. We extend MSPC to call consensus regions across any number of replicated samples, accounting for differences between biological and technical replicates. We observed several master transcription regulators (e.g., SP1 and GATA3) and HDAC2-GATA1 regulatory networks on rescued regions.

**Availability and implementation:** An implementation of the proposed method and the scripts to reproduce the performed analysis are freely available at https://genometric.github.io/MSPC/, MSPC is distributed as a command-line application, an R package available from Bioconductor (https://doi.org/doi:10.18129/B9.bioc.rmspc), and a C# library.

## Introduction

Chromatin immunoprecipitation (ChIP), followed by massively parallel DNA sequencing (ChIP-seq), has become the standard omics technique for the genome-wide localization of *in vivo* DNA-protein binding loci commonly known as “peaks”. A peak is generally inferred when the ChIP-seq read distribution differs significantly from the background signal, and its corresponding *p*-value is more stringent than a given threshold.

In general, peak evaluation is intrinsically limited due to the lack of annotations for “true” binding sites, especially for experimental conditions in which biological signals might be shifted, changed, or depleted [1]. Additionally, peak evaluation is complex, as gene expression regulation involves interactions between combinatorial transcription factor binding sites and chromatin states. Additionally, given the intrinsic noise of the ChIP-seq protocol that leads to artifactual peaks and poor localization of binding loci, peak calling methods are susceptible to high false-positive rates [2–4]. Peak callers utilize various methods to lower the false-positive rate. For instance, MACS [2] uses negative controls such as IgG (non-specific antibody-targeted ChIP-seq), and Ritornello [3] uses peak shape to differentiate between artifactual and “true” binding sites.

A common consensus suggests that “true” binding sites are reproducible across replicated samples (i.e., colocalized within a certain distance), whereas non-overlapping peaks are categorized as either of the following: (a) Characterizing “true” biological variability when studying biological replicates [5]; or (b) Artifactual binding or noise, particularly when studying technical replicates. Accordingly, replicated samples remain a reliable source of information to identify “true” binding sites. However, calling such binding sites across replicated samples lacks gold standards, and it is associated with several open challenges, in particular for studying samples with low variability and high signal-to-noise ratio (technical replicates) and high variability with a low signal-to-noise ratio (biological replicates with heterogeneous cell populations) [6].

MSPC [7,8] and Irreproducible Discovery Rate (IDR [9]) are among the methods used to identify reproducible peaks across replicated samples. IDR uses a copula mixture model to estimate the reproducibility of each pair of peaks in two replicates and to compute the expected rate of irreproducible discoveries [9]. MSPC uses replicates to improve the sensitivity and specificity of peak calling on each sample. MSPC rescues weak peaks; in other words, it differentiates the weak binding sites which are reproducible across replicated samples from background signals (i.e., artifactual binding sites), exploiting the differences between biological and technical replicates. Importantly, it can work with any number of replicates.

The binding sites that are reproducible across all the replicated samples are commonly referred to as *consensus regions*, and both MSPC and IDR can identify them. IDR ranks pairs of peaks in the two replicates based on their irreproducible discovery rate and combines those peaks with rates below a threshold. MSPC first improves the sensitivity and specificity of each replicate and identifies their true-positive peaks using the Benjamini-Hochberg procedure, then identifies consensus regions by merging the true-positive peaks and assigns each a combined stringency score (χ^2^ and right-tail probability).

The present study assesses the biological validity of the peaks MSPC and IDR identify as “true binding sites” and the consensus regions they yield. Accordingly, we developed a feature enrichment test. Our results suggest that MSPC identifies more true binding sites and consensus regions than IDR, encompassing the IDR-identified regions in large. Additionally, our results show that the identified regions are enriched in biologically meaningful annotations and fully encompass essential information needed to understand genomic regulatory networks. For instance, we show the recovery of a large-scale enhancer regulatory network, depending on HDAC2 and GATA1 rescued peaks, whose components are involved in Chronic Myeloid Leukemia (CML) and several cancer-associated processes [10–15].

Identifying “true” binding sites (in noisy samples in particular) and consensus regions have a significant impact on studying high-throughput sequencing data [16,17] with numerous applications spanning from improving sensitivity and specificity of peak callers to studying spatial dependency regulations and combinatorial transcription factor binding in different chromatin states [18,19]. The high-throughput sequencing data are available from public repositories such as ENCODE [20], Roadmap Epigenomics [21], and GEO [22], and are widely adopted for numerous biomedical studies. The quantity and quality of the reproducible regions identified in these samples can profoundly affect any downstream inferences. For instance, the high throughput sequencing data have been used for studying transcription factor regulatory networks [23–27], where identified peaks can vastly influence the topology and connectivity of the regulatory networks, including the inferred causal relationships [1]. Therefore, the results of the present study motivate utilizing methods such as MSPC and IDR to increase the specificity and sensitivity of peak callers and identify consensus regions.

## Materials and methods

In the following, we first provide a brief literature review on the peak callers, we then discuss the characteristics of MSPC and IDR, and finally we define a functional enrichment test to assess the biological validity of the MSPC- and IDR-identified peaks.

### Characteristics of peak callers

A plethora of *peak calling* methods has been developed (reviewed in [1,28–30]). In general, they differ in their statistical model and the number of input signals they operate on.

#### Statistical model

Peak callers identify binding affinities by either scanning the entire genome using a sliding window and test for differential binding between ChIP and control samples at each window based on the Poisson model or its extensions (e.g., MACS [2], PePr [31], and csaw [32]), or using a Hidden Markov Model approach (HMM, e.g., HPeak [33], ODIN [34], histoneHMM [35], and THOR [6]). The sliding window-based methods are sensitive to the window size, where large windows may fail to detect putative peaks (e.g., transcription factor binding sites) while narrow windows may generate severely fragmented peaks on wider binding sites (e.g., histone modifications). In general, methods using HMM can better detect subtle changes as they partition the signal into windows of varying sizes [6].

#### Number of input samples

Concerning the number of input samples, peak callers are generally divided into two groups. First group models binding affinity based on the signal of a single ChIP-seq assay (e.g., MACS). The second group jointly models binding affinities across replicated samples to identify combinatorial enrichment patterns (e.g., [18,19,31,36–38]), they do so either by building models based on single samples then combine them (e.g., jMOSAiCS relies on MOSAiCS [36]), or based on HMM (e.g., [6,35,39]), or sliding window-based approaches (e.g, [31,40–42]). In general, most differential peak calling methods are implemented using the sliding window approach, while a very few HMM-based approaches support replicated samples (e.g., THOR [6]). A possible shortcoming of the HMM-based approaches is that they model a ChIP-seq signal using a limited number of hidden states, which may result in less sensitivity to quantitative changes in signals of closely related conditions [43].

### Characteristics of MSPC and IDR

MSPC [7,8] and IDR [9] are among the methods used to lower false-positive rates and identify consensus regions between replicated samples (see Table 1). IDR measures consistency between two replicates in high-throughput experiments. This quantitative irreproducibility score can then be used to rank pairs of peaks in the two replicates, determine a cutoff for irreproducibility and combine the two replicates. IDR uses a copula mixture model for estimating the expected irreproducible discovery rate of each pair of peaks in two replicates, yielding the expected rate of irreproducible discoveries [9].

**Table 1.** Characteristics of MSPC and IDR.

|  | Model | Multiple Hypothesis Testing Correction | Replicate Count | Output Score |
| --- | --- | --- | --- | --- |
| MSPC | Fisher's combined probability test | False Discovery Rate (FDR, Benjamini-Hochberg procedure) | Unlimited | $\chi^2$, combined $p$-value |
| IDR | Gaussian copula mixture model | Local irreproducible discovery rate (idr) | 2 | Expected irreproducible discovery rate |

Calling consensus regions using IDR falls short in two areas. First, IDR is developed for conservative peak detection, where only highly reproducible peaks across samples are called. Hence, it fails to call peaks in samples with large variance such as biological replicates, where strong peaks on one replicate do not colocalize with peaks from other replicates with a low signal-to-noise ratio (SNR) [1,6]. Low SNR may not only arise due to poor sample quality, rather it can be reflective of true variability between biological replicates, low quantities of starting biological material, or antibody deficiency [44,45]. Second, similar to other methods in this category, it relies on the candidate regions called by the peak caller, hence it may fail to detect subtle changes [17].

MSPC *rescues* weak peaks and identifies consensus regions across any number of replicates; it is perceived to address the aforementioned shortcomings of IDR. MSPC processes biological and technical replicates differently, hence it differentiates between true variability between biological replicates and artifactual binding sites. Therefore, it lowers the false-negative rate between samples with large variance (expected in biological replicates) while preserving a low false-positive rate. To alleviate dependence on the peak caller’s candidate regions, it is suggested to run MSPC on peaks called with a permissive p-value threshold (e.g., 1e-4)[7]. Such a setting would lead to calling a large number of false-positives and a very small number of false-negatives, hence minimizing the probability of missing a true, yet weak binding site. MSPC uses combined stringency of peaks colocalized across replicated samples to differentiate between artifactual and weak binding sites, hence decreasing the number of false-negatives with least false-positives.

For each peak on a sample, MSPC finds the peaks in the other samples overlapping with it. If the number of overlapping peaks is more than a user-defined threshold, it then combines their *p*-values using Fisher’s combined probability test, yielding a combined stringency, χ^2^, and the corresponding combined p-value. MSPC *confirms* the overlapping peaks if the combined χ^2^ is larger than a user-defined threshold, and *discards* if otherwise. A peak might be tested multiple times if it overlaps with multiple peaks on another sample. Therefore, a peak might be *confirmed* based on one test and *discarded* based on another. When samples are biological replicates, MSPC *confirms* a peak if it passes at least one test (heterogeneity may reflect true biological variability), and with technical replicates, MSPC *discards* a peak if it does not pass all the tests (since more homogeneity is expected in this case). The confirmed peaks are then corrected for false-discovery rate using the Benjamini–Hochberg procedure.

MSPC calls a consensus region where true-positive peaks on either of the replicates suggest binding loci. The coordinates of a consensus region are the union of overlapping true-positive peaks across all the samples, and its stringency is determined by combining the *p*-values of the overlapping peaks using the Fisher’s combined probability test (see Supplementary Figure 1).

### Data pre-processing

#### ENCODE data preprocessing

ChIP-seq raw data were downloaded from ENCODE (see Supplementary Table 1), peaks on each sample were called using MACS2 with the following options: --*mfold 5, --bw 300 --pvalue 0*.*0001*. For each sample we used control samples as linked on ENCODE for each experiment; some experiments use a common control between multiple replicates (e.g., https://www.encodeproject.org/experiments/ENCSR532KTI/), some experiments use different control samples for each replicate (e.g., https://www.encodeproject.org/experiments/ENCSR121PFY/), or use two controls for each sample (e.g., https://www.encodeproject.org/experiments/ENCSR574XEO/).

#### Genomic annotations and optimal MSPC threshold set

Our functional enrichment procedure was applied to a set of 50 randomly chosen ENCODE transcription factors (TFs) as listed in the Supplementary Table 1. For each TF, the procedure was repeated using 28 different sets of MSPC thresholds (see Supplementary Table 2). The threshold sets were chosen from conservative to highly permissive, in order to cover a broad number of possible pools of rescued peaks. The best thresholds were defined as the ones producing the highest enrichment score (i.e., the highest z-score of the enrichment test, see details in the next Section) for each TF. The threshold set -*w 1E-04* (weak significance threshold), *-s 1E-08* (stringent significance threshold), and -*g 1E-06* (combined significance threshold), was the one yielding the best enrichment score for every TF. To evaluate the enrichment for each TF, we selected 9 genomic annotations (genome assembly hg38), whose loci were downloaded from the UCSC Genome Browser database ([46] accessed on 2020-01-31): CpG islands, DNase clusters, enhancers, exons, introns, promoters, coding RefSeq genes, noncoding RefSeq genes, and non-RefSeq transcripts. We chose these annotations to have a straightforward measure of peak enrichment at open chromatin regions (DNase clusters), transcripts (exons, introns, coding RefSeq, noncoding RefSeq, and non-RefSeq transcripts), and both proximal and distal regulatory elements (promoters, CpG islands, enhancers).

### Functional enrichment test

The objective of the validation procedure is to assert if MSPC-rescued peaks are enriched in biologically meaningful loci. Accordingly, we defined three types of genomic regions: peaks, annotations, and regions not covered by any of them; where the first two may overlap to some extent. The higher the overlap between peaks and annotations, the higher the ability of the peak caller/rescuer to recall functional genomic regions. Since the number and coverage of a specific annotation is fixed for a given database, the only variables we need to consider are the number and position of called peaks. In particular, we define the conditional probability of a nucleotide overlapping a peak to contain an annotation, and the conditional probability of a nucleotide non-overlapping a peak to contain an annotation (see Supplementary Material for details). The difference *β* = − measures the ability of the peak caller/rescuer to recall functional annotations, such that if *β* > 0 (i.e., >), there is a higher probability of observing a peak at a random position within an annotated region. Therefore, the greater the *β* value, the higher the genome-wide proportion of annotated nucleotides within peaks. Our objective is to assert if the MSPC-rescued peaks on a given sample are strongly enriched in functional annotations w.r.t a standard baseline peak set given by IDR consensus on the same sample.

For both MSPC-rescued and IDR consensus peaks, we computed the enrichment score as the ratio = (−_0_)/, where *β*_0_ = 0 (i.e., =) is the *β* value under the null hypothesis, and is the standard error, that is the standard deviation of the sampling distribution of *β*, assuming that the underlying distribution of under the null hypothesis is well approximated by a gaussian distribution with mean 0 and standard deviation equal to 1 (see Supplementary Material for further details). In this way, we can both assess the significance of annotation enrichment and directly compare the enrichment scores for MSPC-rescued peaks against IDR consensus, for each annotation and sample (i.e., transcription factor). The scripts for the functional enrichment test are freely available from https://github.com/Genometric/MSPC/tree/dev/ValidationScripts.

### Overrepresentation analysis and motif search

TF binding motif enrichment was performed using MEME-ChIP with default settings [47] available at https://web.mit.edu/meme_v4.11.4/share/doc/meme-chip.html. Motif enrichment was evaluated using the threshold E-value < 1E-10.

Overrepresentation analysis against the KEGG pathways and ChEA TF databases was done using the Enrichr online tool [48] available at https://maayanlab.cloud/Enrichr.

### HDAC2-GATA1 enhancer regulatory network reconstruction

For each of the 48 TFs considered in this study, we obtained the list of TFB motifs enriched at MSPC rescued enhancers (enrichment E-value < 1E-10; Supplementary Table 1). HDAC2 was the TF showing the strongest motif enrichment for another TF in the set of 48: GATA1. This means that HDAC2 rescued peaks at known enhancers are enriched in GATA1 binding motifs and, therefore, these enhancers are common HDAC2-GATA1 targets. Moreover, since the enrichment in GATA1 motifs is exactly at HDAC2 rescued peaks, common target enhancers should bind HDAC2-GATA1 simultaneously.

To further investigate the impact of this rescued regulatory network, we considered HDAC2 rescued enhancer peaks overlapping GATA1 rescued enhancer peaks, and considered the set of closest TSS within 100 kb from these peaks, referred to as the HDAC2-GATA1 target gene set. We chose a 100 kb maximum distance to reduce enhancer-TSS false-positive associations, in accordance with recent literature [27,55]. To evaluate the importance of the rescued HDAC2-GATA1 targets, we performed overrepresentation analysis (ORA) with three goals: (i) assess disease and pathway enrichment of the HDAC2-GATA1 target genes through the KEGG database (Supplementary Table 2; [51]); (ii) check if HDAC2-GATA1 target genes are enriched in transcriptional master regulators, and (iii) evaluate if these genes are regulatory targets in specific cell lines [52,57]. We performed ORA using the online enrichment analysis tool Enrichr [58].

## Results and discussion

### MSPC enrichment-based assessment

To have a reference set of reproducible peaks for each transcription factor (TF), we run IDR 2.0.4 (available at https://github.com/nboley/idr) with a global IDR threshold of 0.05 from the output consensus regions. This assessment has two goals: (i) Verify the number of reference IDR peaks that are also detected by MSPC, and (ii) Assert if those MSPC-rescued peaks that are not in the IDR set are enriched in functionally important genomic regions (i.e., annotations). Notably, for every TF, MSPC was able to find all the reproducible IDR peaks; we name them as the “common” set of peaks. The scripts to reproduce the performed analysis are available from https://github.com/Genometric/MSPC/tree/dev/ValidationScripts.

Rescued MSPC peaks not present in the common peak set are referred to as *MSPC-specific peaks set*. To assess the biological relevance of specific peaks, we compared their functional enrichment score (i.e., the z score of the enrichment test) to the score of the common set of peaks (Supplementary Figure 2). For every annotation, MSPC-specific peaks showed higher enrichment than common ones (Figure 1). Accordingly, MSPC confirms reproducible peaks and rescues peaks whose functional role is not negligible. Notably, the most significant MSPC enrichments against common peaks were at enhancers, promoters, and DNase clusters, denoting MSPC’s best performances in rescuing critical regulatory and accessible chromatin regions. At the TF level, MSPC showed higher enrichment in 45/48 TFs with respect to common peaks, meaning that it rescues critical genome-wide TF enrichments that would be lost otherwise.

**Figure 1.**
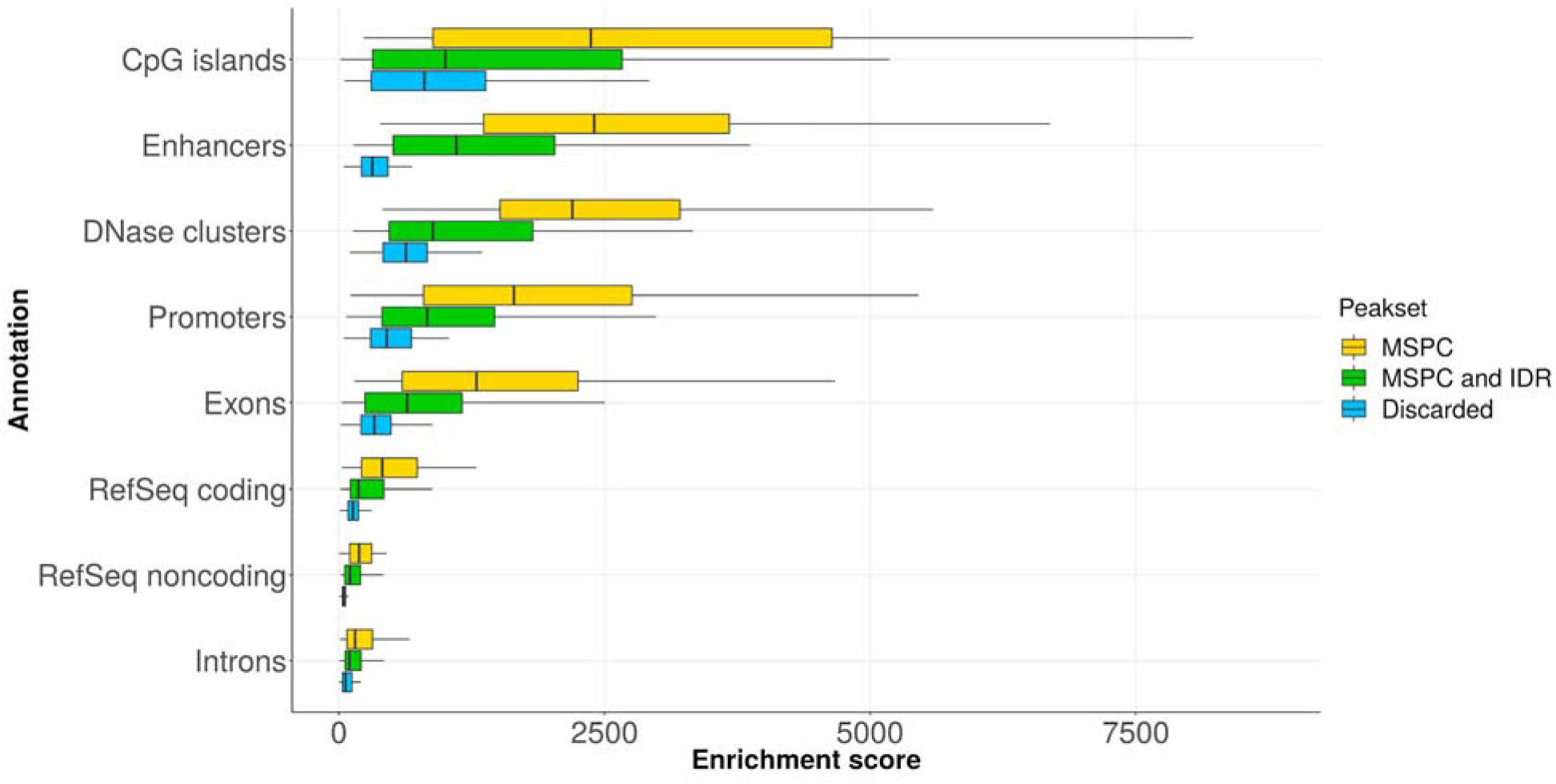
Enrichment score distribution (x axis) for MSPC discarded peaks (cyan box), MSPC rescued peaks discarded by IDR (i.e., MSPC-specific peaks set; yellow box), and peaks retained by both MSPC and IDR (i.e., common peaks; green box), aggregated by 9 hg38 annotations (y axis). For each of the 48 TFs analyzed, there were no peaks retained by IDR and discarded by MSPC (i.e., MSPC always included IDR results). The 9 hg38 annotations include: CpG islands (n = 31,144), Enhancers (n = 393,964), DNase clusters (n = 2,107,358), Promoters (n = 34,996), Exons (n = 313,276), RefSeq coding transcripts (n = 67,635), RefSeq non-coding transcripts (n = 17,271), Introns (n = 172,751). Coordinates for hg38 annotations were downloaded from the UCSC Genome Browser (accessed on: 2020-01-31) [46].

### Motif enrichment at rescued enhancers

MEME ChIP transcription factor binding (TFB) motif enrichment at rescued enhancers shows the presence of several transcription master regulators. The most frequent enriched motif is GATA3 (20/48 TFs), which has been recently described as a key factor in enhancer-dependent cell reprogramming [11] and T-cell differentiation [12]. The second most frequent motif (19/48 TFs) is SP1, known for binding enhancers, regulating chromatin looping [49] and playing a key role in malignant hematopoiesis, through its interaction with GATA1 [10]. The critical role of chromatin looping is also demonstrated by the occurrence of CTCF binding site enrichments (the third-most enriched motif, with 18/48 TFs). CTCF is a widely studied insulator which is recognized as one of the main designers of topologically associated domains (TADs). TADs insulate portions of active chromatin, determining how and when genomic DNA is processed (e.g., transcribed and/or replicated). Although TADs are conserved among evolutionary-related species, cancer-associated cell fate reprogramming is often associated with mutated TAD boundaries [50]. Other cancer-associated TFB motif enrichments have been found as well, including RUNX1-RUNX3 involved in lymphoid cell differentiation [13,14], and ETV6, involved in lymphoid malignant transformation [15]. The complete table of TFB motif enrichments at rescued enhancers is reported in Supplementary Table 3.

### HDAC2-GATA1 rescued regulatory network

We identified 26,514 HDAC2 and 2,513 GATA1 enhancers (see Methods and the Supplementary methods section: “TFB motif enrichment at enhancers”) at MSPC-rescued binding loci (after removing peaks shorter than 200 bp); among them, 1,627 peaks are in the *common set*. Over-representation analysis against the KEGG database (see Supplementary Table 4; [51]) confirmed Chronic Myeloid Leukemia (CML) as the most enriched pathway (adjusted *p*-value = 1.37E-03), with 25 CML genes as targets of rescued enhancers. Other leukemia-related pathways were significantly enriched, including: VEGF signaling (19 target genes; adjusted *p*-value = 5.19E-03), Calcium reabsorption (17 target genes; adjusted *p*-value = 5.23E-03), Cell cycle (31 target genes; adjusted *p*-value = 5.26E-03), Rap1 signaling (43 target genes; adjusted *p*-value = 0.0186), Cellular senescence (35 target genes; adjusted *p*-value = 0.0201), Platelet activation (28 target genes; adjusted *p*-value = 0.0313), Leukocyte transendothelial migration (26 target genes; adjusted *p*-value = 0.0314). In addition, GATA1 and GATA2 binding in K562 cells were the top overrepresented terms (adjusted p-value: 9.04E-97 and 1.61E-86, respectively; Supplementary Table 5) among target genes, against the ChEA TF database [52], indicating how both rescued enhancers and their associated genes are targets of GATA1 regulatory network. Collectively, these results show how MSPC may successfully recover genome-wide enrichments (i.e., peaks) that are part of the K562 CML regulatory networks, coherently with the sample cell line and phenotype.

## Conclusion

We argue the significant impact of improving the sensitivity and specificity while identifying binding affinities on high-throughput sequencing data by discussing the biological characteristics unveiled using weak but reproducible binding sites. Specifically we show how these rescued peaks are enriched in biologically meaningful information. This information emerges from the overrepresentation of genomic elements, including promoters, CpG islands, enhancers, and DNase clusters (regions of open chromatin), suggesting that these weak but reproducible elements are part of large-scale active chromatin networks (e.g., active enhancers and transcribed genes). We showed how one of these largest rescued regulatory networks is represented by the enhancers enriched in HDAC2-GATA1 peaks, which neighbouring genes are involved in chronic myeloid leukemia-associated processes and K562-specific regulation.

We discuss two methods for differentiating between weak and artifactual binding sites and calling consensus regions across replicated samples: MSPC and IDR. Our analysis over K562 ENCODE data showed that MSPC contains all IDR-identified reproducible regions, in addition to “rescuing” many other biologically relevant weak regions. Additionally, MSPC consensus regions that are not common to IDR show a larger enrichment by annotation (8/8 genomic annotations; Figure 1) and by TF (45/48 TFs; Supplementary Figure 2). Accordingly, the consensus regions identified by MSPC provide a more appropriate set of informative genomic regions, favouring discovery over conservativeness, while controlling false positives. Since MSPC is applied at post-peak calling, it can be used to produce a single set of peaks from multiple replicates, as well as from multiple sets of peaks obtained by applying different peak calling methods [53].

Both MSPC and IDR operate on regions called using peak callers. Hence, their candidate sites are limited to the regions identified by the peak caller. To alleviate this limitation, a recommended practice for MSPC is to call peaks using a permissive *p*-value threshold to minimize the probability of missing weak binding sites at the cost of increasing false-positive rate; our assessment shows that MSPC can distinguish between true weak binding sites and artifactual regions in an input with a high false-positive rate. Additionally, given that the statistical model of both methods rely on regions binding affinity, they have limited application in sequencing protocols where there is not sufficient evidence to reason about the statistical significance of binding affinity (e.g., single-cell protocols such as ATAC-seq).

