## Supplementary Material for "Rescuing Biologically Relevant Consensus Regions Across Replicated Samples"

**Supplementary Table 1.** Summary of the experiments used in this paper. The experiments are randomly selected from the immortalized K562 cell line available from ENCODE.

| Index | ENCODE Experiment ID | Target Gene | Replicate Count | Control Count |
| --- | --- | --- | --- | --- |
| 1 | ENCSR000BLR | SIN3A | 2 | 1 |
| 2 | ENCSR000BQY | PML | 2 | 1 |
| 3 | ENCSR000EFR | ZC3H11A | 2 | 1 |
| 4 | ENCSR000EGD | BACH1 | 2 | 1 |
| 5 | ENCSR000EGJ | MYC | 2 | 1 |
| 6 | ENCSR000EGL | IRF1 | 2 | 1 |
| 7 | ENCSR000EGX | MAFK | 2 | 1 |
| 8 | ENCSR000EWF | YY1 | 2 | 1 |
| 9 | ENCSR000EWM | GATA1 | 2 | 1 |
| 10 | ENCSR000FCE | ETV6 | 2 | 2 |
| 11 | ENCSR030TJP | DACH1 | 2 | 2 |
| 12 | ENCSR051DXE | FUS | 2 | 2 |
| 13 | ENCSR075HTM | HDAC2 | 2 | 2 |
| 14 | ENCSR086FZL | KAT8 | 2 | 1 |
| 15 | ENCSR112RNT | HNRNPLL | 2 | 2 |
| 16 | ENCSR121PFY | CDC5L | 2 | 2 |
| 17 | ENCSR154EIH | TRIP13 | 2 | 2 |
| 18 | ENCSR167KBO | ZNF316 | 2 | 2 |
| 19 | ENCSR175SZH | ZSCAN29 | 2 | 2 |
| 20 | ENCSR177DNR | FIP1L1 | 2 | 2 |
| 21 | ENCSR189PYJ | SMAD2 | 2 | 2 |
| 22 | ENCSR189TRZ | TCF12 | 2 | 2 |
| 23 | ENCSR213HBY | TOE1 | 2 | 2 |
| 24 | ENCSR249BHQ | ZNF592 | 2 | 2 |
| 25 | ENCSR264CZJ | THRA | 2 | 2 |
| 26 | ENCSR286PCG | ZBED1 | 2 | 2 |
| 27 | ENCSR334HSW | ZNF318 | 2 | 2 |
| 28 | ENCSR414TYY | RUNX1 | 2 | 2 |
| 29 | ENCSR426URK | AFF1 | 2 | 2 |
| 30 | ENCSR446LAV | DDX20 | 2 | 2 |
| 31 | ENCSR512NLO | MNT | 2 | 2 |
| 32 | ENCSR532KTI | GTF2E2 | 2 | 1 |
| 33 | ENCSR547LKC | GATAD2B | 2 | 2 |
| 34 | ENCSR574XEO | NUFIP1 | 2 | 2 |
| 35 | ENCSR598GER | CCAR2 | 2 | 2 |
| 36 | ENCSR657JLK | SIN3B | 2 | 2 |
| 37 | ENCSR664AOA | TRIM25 | 2 | 2 |
| 38 | ENCSR675LRO | MLLT1 | 2 | 2 |
| 39 | ENCSR686EYO | KHSRP | 2 | 2 |
| 40 | ENCSR731LHZ | E4F1 | 2 | 2 |
| 41 | ENCSR757IIU | HMBOX1 | 2 | 2 |
| 42 | ENCSR815ZDS | SREBF1 | 2 | 2 |
| 43 | ENCSR871TKJ | THRAP3 | 2 | 2 |
| 44 | ENCSR908CMW | KDM1A | 2 | 2 |
| 45 | ENCSR914NEI | MTA3 | 2 | 2 |
| 46 | ENCSR931HNY | NCOA1 | 2 | 2 |
| 47 | ENCSR987PBI | DNMT1 | 2 | 2 |
| 48 | ENCSR998AJK | NRF1 | 2 | 2 |

**Supplementary Table 2.** The sets of MSPC thresholds used in the experiments.

|  | Weak Significance Threshold (-w) | Stringent Significance Threshold (-s) | Combined Stringency Threshold (-g) |
| --- | --- | --- | --- |
| 1 | 1.00E-01 | 1.00E-02 | 1.00E-02 |
| 2 | 1.00E-01 | 1.00E-03 | 1.00E-02 |
| 3 | 1.00E-01 | 1.00E-03 | 1.00E-03 |
| 4 | 1.00E-01 | 1.00E-04 | 1.00E-03 |
| 5 | 1.00E-01 | 1.00E-04 | 1.00E-04 |
| 6 | 1.00E-02 | 1.00E-04 | 1.00E-04 |
| 7 | 1.00E-02 | 1.00E-05 | 1.00E-02 |
| 8 | 1.00E-02 | 1.00E-05 | 1.00E-03 |
| 9 | 1.00E-02 | 1.00E-05 | 1.00E-04 |
| 10 | 1.00E-03 | 1.00E-05 | 1.00E-05 |
| 11 | 1.00E-03 | 1.00E-06 | 1.00E-05 |
| 12 | 1.00E-03 | 1.00E-06 | 1.00E-06 |
| 13 | 1.00E-03 | 1.00E-07 | 1.00E-05 |
| 14 | 1.00E-03 | 1.00E-07 | 1.00E-06 |
| 15 | 1.00E-03 | 1.00E-07 | 1.00E-07 |
| 16 | 1.00E-03 | 1.00E-08 | 1.00E-04 |
| 17 | 1.00E-03 | 1.00E-08 | 1.00E-05 |
| 18 | 1.00E-03 | 1.00E-08 | 1.00E-06 |
| 19 | 1.00E-03 | 1.00E-08 | 1.00E-07 |
| 20 | 1.00E-03 | 1.00E-08 | 1.00E-08 |
| 21 | 1.00E-04 | 1.00E-05 | 1.00E-05 |
| 22 | 1.00E-04 | 1.00E-06 | 1.00E-05 |
| 23 | 1.00E-04 | 1.00E-06 | 1.00E-06 |
| 24 | 1.00E-04 | 1.00E-07 | 1.00E-07 |
| 25 | 1.00E-04 | 1.00E-08 | 1.00E-08 |
| 26 | 1.00E-05 | 1.00E-06 | 1.00E-06 |
| 27 | 1.00E-05 | 1.00E-07 | 1.00E-07 |
| 28 | 1.00E-05 | 1.00E-08 | 1.00E-08 |

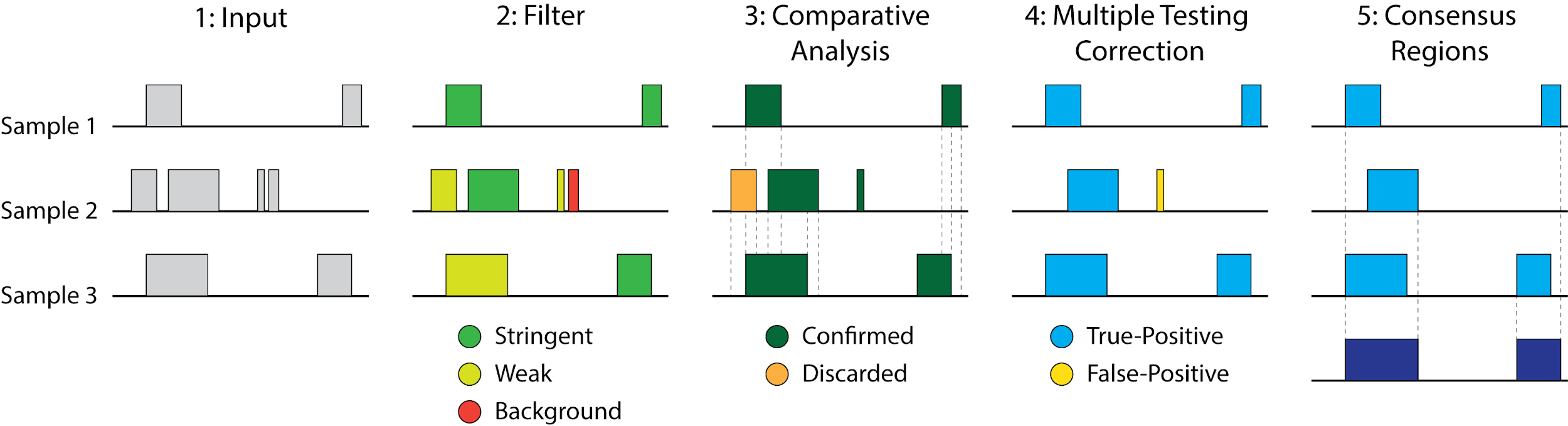

**Supplementary Figure 1.** An illustration of processing regions on three replicated samples using MSPC.

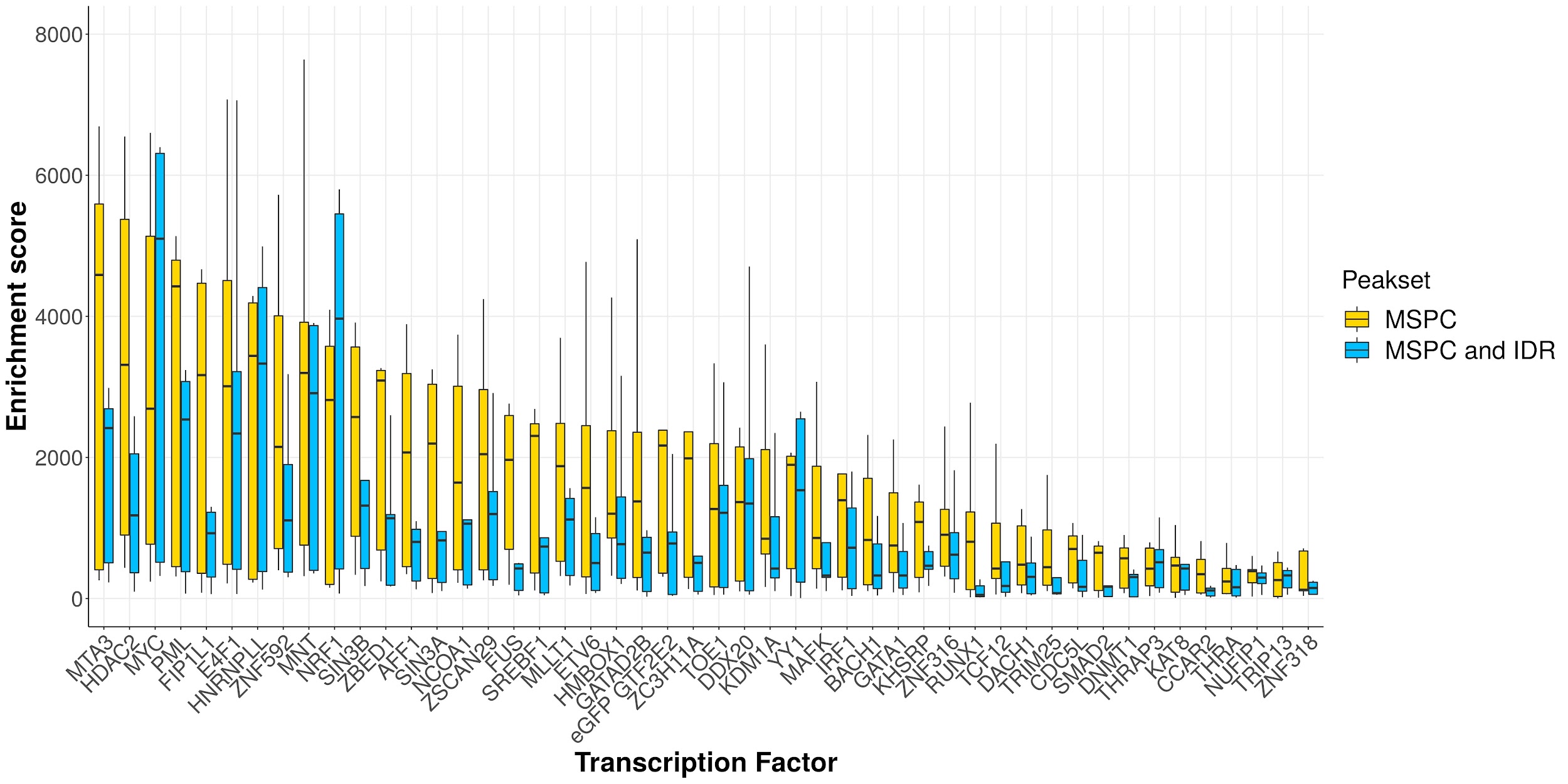

**Supplementary Figure 2**. Enrichment score distribution (y-axis) for peaks retained by both MSPC and IDR (i.e., common peaks; cyan boxes) and rescued by MSPC but discarded by IDR (i.e., MSPC-specific peaks set; yellow boxes), aggregated by 48 ENCODE TFs (x-axis). Among the 48 TFs analyzed, 44 showed higher median genomic annotation enrichments for MSPC-specific peaks (yellow boxes). The remaining 4/48 TFs (the first three TFs on the left) showed significant enrichment in genomic annotations (yellow boxes), although not higher than common peaks (cyan boxes).

**Supplementary Table 3**. This table shows the occurrence of transcription factor binding motifs (TFBMs) within MSPC-rescued enhancers associated with the 48 TFs. For each TF, a motif is considered to be enriched within enhancers and added to the table if the E-value reported by MEME-ChIP (either MEME or DREME algorithm) is below 1E-04.

**Supplementary Table 4**. KEGG pathways enrichment. KEGG overrepresentation analysis was done using the Enrichr suite (<https://maayanlab.cloud/Enrichr/>) using the list of genes in the neighborhood (< 100 kb) of HDAC2-GATA1 associated enhancers. For each KEGG pathway, the following information is shown: number of genes of the list belonging to the pathway, P-value, Benjamini-Hochberg adjusted P-value, enrichment odds ratio, Enrichr combined score, symbols of genes in the list that belong to the pathway.

**Supplementary Table 5**. ChEA transcription factor enrichment. ChEA overrepresentation analysis was done using the Enrichr suite (<https://maayanlab.cloud/Enrichr/>) using the list of genes in the neighborhood (< 100 kb) of HDAC2-GATA1 associated enhancers. For each TF in ChEA, the following information is shown: number of genes of the list belonging to the pathway, P-value, Benjamini-Hochberg adjusted P-value, enrichment odds ratio, Enrichr combined score, symbols of genes in the list that belong to the pathway.

### 1. Modeling of genomic annotations and peaks distribution

Let *G* denote genome; the null probability $P(A)$ of finding a functional annotation by randomly picking a single position in the genome can be estimated as $A/G$, where $A$ is the size in bp (i.e. the genomic coverage) of the annotated portion of the genome. We define the binary random variable $Y\in\left\{ 0,1 \right\}$ to indicate the annotations, i.e. we define $Y=1$ if a nucleotide contains annotation and $Y=0$ if the nucleotide lies outside of annotations. Similarly, we define the binary random variable $X\in\left\{ 0,1 \right\}$ to indicate the nucleotides overlapping peaks found by the peak caller/rescuer. In particular, $X=1$ if the nucleotide overlaps a peak, and $X=0$ if it doesn’t, and we indicate with $U$ the total size of peaks in bp (i.e. the coverage of the union of all peaks).

We then consider the conditional probability of $Y$given $X$, i.e. the probability of a nucleotide to contain an annotation, conditionally of overlapping or non overlapping a peak. In particular, we define $p=P(Y=1 | X=1)$ the probability of a nucleotide overlapping a peak to contain an annotation, and $a=P(Y=1 | X=0)$ the probability of a nucleotide non-overlapping a peak to contain an annotation. We are interested in comparing $p$ and $a$. We estimate the former with $\hat{p}=A_{1}/U$, where $A_{1}$ is the coverage of annotations within peaks (i.e. for which $X=1$) and $U$ is the coverage of the union of all peaks. Similarly, we estimate the latter with $\hat{a}=A_{0}/(G-U)$, where $A_{0}$ is the coverage of annotations outside peaks (i.e. for which $X=0$) and $G-U$ is the amount of genome non-overlapping any peak.

Functional enrichment is expressed as the peak caller/rescuer capability of finding as many annotations as possible with the smallest possible number of peak calls. A large functional enrichment is achieved when $p=(Y=1 | X=1)$ is higher than $a=(Y=1 | X=0)$, therefore we model it as the proportion difference: $\beta=p-a$.

#### Functional Enrichment

Functional enrichment ($\beta$) is defined as the difference between the proportion of the genome converted by peaks that overlap annotations (*p*) and the proportion of the genome without peaks that overlap annotations (*a*):

$$\beta= p - a = P(Y=1 | X=1) - P(Y=1 | X=0)$$

Estimates:

$$\hat{\beta}= \hat{p} - \hat{a} \hat{p}= \frac{A_{1}}{U} \hat{a}=\frac{A_{0}}{G-U}$$

Where G is the genome size (bp count), U is the coverage of peaks (in bp), $A_{0}$is the coverage of annotations outside peaks (in bp), and $A_{1}$is the coverage of annotations within peaks (in bp).

#### Functional enrichment test

We test functional enrichment under the null hypothesis $H_{0}:\beta=0$**,** versus the alternative $H_{1}:\beta\neq0$. Note that, although we are generally interested in testing for positive enrichment (i.e. $\beta>0$) for a given genomic feature, there are no strong assumptions allowing us to employ a one-sided test. For instance, under specific experimental conditions that may cause genome-wide repression of certain genomic elements, the peak caller/rescuer may find only a small number of active genomic regions (i.e., few peaks) and show feature under-representation (i.e.,$\beta<0$). Significant feature under-representation could also arise within difficult genomic regions, such as repetitive or low complexity DNA, where it is hard to obtain reliable estimates of the signal-to-noise ratio and hence of peak presence.

Enrichment significance depends on the estimated effect size, i.e. on the difference $\beta-\beta_{0}$where $\beta_{0}$ is the null enrichment being tested (in our case, ${\beta_{0}=0}$), and on its standard error ($\sigma$). The standard error measures the reliability of the estimated effect size, depending on the proportions we are comparing and the sample size. Under the null hypothesis $H_{0}:\beta=0$ we have $p=a$, so we can compute the pooled sample proportion as $\hat{q}=\left( \hat{p}U+\hat{a}\left( G-U \right) \right)/\left( U+\left( G-U \right) \right)=A/G$and employ it to estimate $\sigma$ as $\hat{\sigma}=\sqrt{\hat{q}\left( 1-\hat{q} \right)\left( 1/U+1/\left( G-U \right) \right)}$. With constant $U$, as $\hat{q}$ approaches 0.5 the standard error is maximized. Conversely, if either $\hat{q}$ or $1-\hat{q}$ is a very small value, $\hat{q}(1-\hat{q})$ is small and so is the standard error. In our case, when the number of annotations is large (i.e. $A$ is large) we obtain a small $\hat{\sigma}$. This happens because a peak corresponding to a regulatory (e.g., histone marker) or processing (e.g., RNA Polymerase II binding) signal, thus being neither a technical artifact nor random background fluctuation, is generally associated with a genomic region having a functional role (either active or repressed, depending on the signal type), that is more likely found within an annotation. Therefore, we expect a larger fraction of enriched peaks in a frequent genomic annotation (e.g., promoters), rather than a rare one (e.g., snoRNA), for which we expect higher $a$ values.

The test statistic is $z=\left( \hat{\beta}-{\beta_{0}} \right)/\hat{\sigma}$, where ${\beta_{0}}$ is the effect size under $H_{0}$ (in our case, ${\beta_{0}=0}$). Under the null hypothesis, $z$ is approximately distributed as a standard Normal $N\left( 0,1 \right)$, hence we can compute the *p*-value as the probability of observing a value as extreme as or more extreme than $\left| z \right|$in a standard Normal distribution, and we can compute the 95% confidence interval for $\beta$ as ${CI}_{95\%}=\left( \hat{\beta}-1.96 \hat{\sigma}; \hat{\beta}+1.96 \hat{\sigma} \right)$. The ${CI}_{95\%}$ containing $\beta=0$ (i.e., the null enrichment) is a synonym of non-significant enrichment at level 5%.

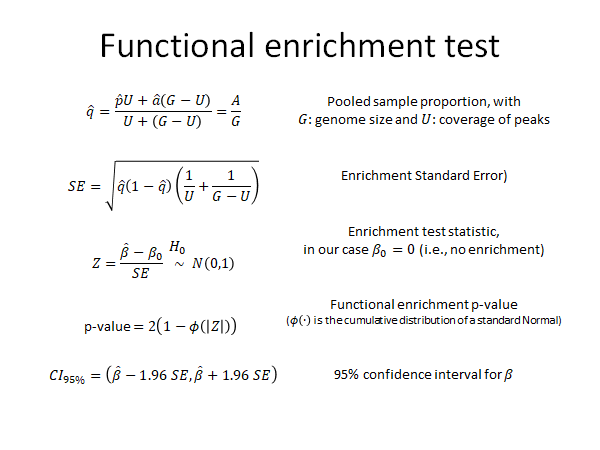

### 2. TFB motif enrichment analysis at enhancers

To further evaluate the biological impact of MSPC rescued peaks, we assessed the presence of regulatory motifs at rescued enhancers. To ensure no overlap between MSPC and IDR results, we considered genomic features overlapping peaks rescued only by MSPC. We chose enhancers for two reasons. Firstly, enhancers have a clear and well-studied impact on gene expression, largely documented on biological knowledge bases, including Gene Cards [[54]](https://paperpile.com/c/hcmzyF/HhZ0F) and the UCSC Genome Browser [[46]](https://paperpile.com/c/hcmzyF/QEUfi), and providing a direct measure of the regulatory importance of rescued peaks. Secondly, part of an enhancer target gene set can be recovered by proximity to the closest TSSs [[55]](https://paperpile.com/c/hcmzyF/b8Mw5), enabling a partial and yet robust evaluation of its impact on genomic regulation. To achieve this goal, we first isolated all MSPC rescued peaks overlapping at least one enhancer, from the hg38 GeneHancer set [[54]](https://paperpile.com/c/hcmzyF/HhZ0F), downloaded from the UCSC Genome Browser (accessed on 2020-01-31). To improve the motif search, we excluded peaks shorter than 200 bp. The remaining peaks were used as input for transcription factor binding (TFB) motif search, using the MEME ChIP suite (see Supplementary File 1 for details; [[56]](https://paperpile.com/c/hcmzyF/s4hNS)).
